# An angiopoietin-2 vaccine improves arteriovenous malformation pathology in hereditary hemorrhagic telangiectasia mice

**DOI:** 10.1101/2025.10.13.682178

**Authors:** Sima Qutaina, Haitian Zhao, Zhiming Wang, Caterina Ivaldo, Santiago Ruiz, Philippe Marambaud

## Abstract

Hereditary hemorrhagic telangiectasia (HHT) is a genetic vascular disorder that causes systemic arteriovenous malformations (AVMs) associated with severe complications. Angiopoietin (ANG)-2 has been identified as a consistently upregulated secreted protein across various HHT models, and neutralizing ANG2 reduces AVMs in mice. ANG2 has thus emerged as a potential target for HHT treatment. Here, we report the development of a peptide vaccine (ANG2-P3:CRM197) that selectively targets ANG2 over ANG1 and tested its effectiveness in decreasing retinal AVMs in neonatal mice injected with BMP9/10 blocking antibodies, a model of HHT. Litter groups from female C57BL/6 mice immunized with ANG2-P3:CRM197 received injections of anti-BMP9/10 antibodies, and their retinas were examined for vascular pathology. The potential toxicity of the vaccine was evaluated in females 12 months post-immunization through echocardiography, basic metabolic panels, and lipid profiles. Circulating anti-ANG2 antibodies were detected in nursing neonates of vaccinated females, with antibody levels comparable between litters and their dams, indicating effective antibody transfer from the dams. A significant decrease in AVM number and size was observed in the retinas of pups exposed to ANG2-P3:CRM197 antibodies compared to unexposed pups. Arterial and venous diameters were normalized in the vaccinated pups’ retinas. The vaccinated females showed no abnormalities in cardiac, liver, or kidney functions. A vaccine strategy targeting ANG2 appears safe and improves AVM pathology in HHT mice. These findings further support the potential of inhibiting ANG2 as a viable approach for treating AVMs in HHT.

**KEY POINTS:**

- A peptide vaccine that targets ANG2 reduced AVM number and size, and normalized arterial and venous diameters in a mouse model of HHT.
- The vaccine did not cause any abnormalities in cardiac, liver, or kidney functions, and thus appeared safe.

## INTRODUCTION

Hereditary hemorrhagic telangiectasia (HHT), an autosomal dominant genetic disorder affecting between 2 and 12 in 5,000 individuals, is characterized by the focal development of arteriovenous malformations (AVMs) in multiple tissues^1,2^. AVMs are fragile, enlarged blood vessels that form high-flow shunts between arteries and veins. These malformations affect various tissues and can lead to severe complications, including heart failure, internal bleeding, and anemia, which result in a poor quality of life and make HHT a resource-intensive and costly disease^3–6^. Loss-of-function (LoF) heterozygous germline mutations associated with HHT are found in the genes *ENG* (which encodes endoglin; HHT type 1 (HHT1)), *ACVRL1* (which encodes activin receptor-like kinase 1, ALK1; HHT type 2 (HHT2)), and *SMAD4* (which encodes mothers against decapentaplegic homolog 4, Smad4; juvenile polyposis-HHT combined syndrome)^7^. ALK1, ENG, and Smad4 all participate in the same TGF-β signaling pathway in endothelial cells (ECs). ALK1 is a bone morphogenetic protein (BMP) type I receptor that forms complexes with a BMP type II receptor and the co-receptor ENG to bind circulating ligands BMP9 and BMP10. Activation of the ALK1 receptor, through Smad1/5-Smad4 signaling, regulates gene expression programs involved in vascular development and maintenance^8^.

Downstream of ALK1 signaling disruption, the molecular changes that lead to AVM development in HHT are complex and not fully understood. Studies have aimed to uncover transcriptional signatures in HHT mouse models to fill this knowledge gap. Data from our group and others have demonstrated that the angiogenic growth factor angiopoietin-2 (ANG2)^9,10^ is upregulated in various HHT mouse models. RNAseq and qPCR analyses revealed an increase in *Angpt2* mRNAs (which encode ANG2) as the most significant gene expression change in retinal ECs of neonates treated with BMP9 and BMP10 neutralizing antibodies (BMP9/10ib)^11^, a model of HHT. Further RNAseq and CHIPseq analyses showed that Smad4 directly represses *Angpt2* transcription in ECs, and that ANG2 inhibition through the neutralization antibody LC10 prevents AVMs and normalizes vessels in the retina of Smad4 inducible EC-specific knockout (iECKO) mice^12^, another HHT model. More recently, the Meadows team used RNAseq analyses to demonstrate that a universal increase in endothelial ANG2 expression is present in the brains of *Acvrl1, Eng*, and *Smad4* iECKO mice. They also found that ANG2 blockade with LC10 improved brain vascular pathologies across all three models^13^. These findings align with previous studies demonstrating elevated ANG2 expression in several other *Acvrl1-*deficient models^14,15^, highlighting the central role of ANG2 in HHT pathogenesis. Here, we describe the development of a vaccine targeting ANG2 and examine its effects on retinal AVM development in BMP9/10ib mice.

## METHODS

### Vaccine preparation and mouse injection

The selected ANG2 epitopes (ANG2-P1, -P2, and -P3; GenScript) were conjugated to the CRM197 carrier protein as described in the established protocol^16^ with minor modifications. The epitopes were chemically coupled with the carrier by extending each peptide with a short linker region containing two glycine residues and an N-terminal cysteine. The peptides were dissolved in DMSO at 4 mg/mL, while CRM197 (VCar-Lsx004, Creative Biolabs) was prepared in Conjugation buffer [PBS (#28372, Pierce), 2mM EDTA] at 1 mg/mL. To activate CRM197, a 26-fold molar excess (1:10 dilution) of succinimidyl-4-(N-maleimidomethyl) cyclohexane-1-carboxylate (SMCC) cross-linker (#A35394, Pierce) was added to a concentration of 1.5 mg/mL in DMSO. The mixture was incubated at room temperature (RT) for 30 min. Following this, desalting was performed using a resin column (Zeba Desalt Spin Column, Pierce) to remove unreacted crosslinker and byproducts. The activated CRM197 volume was adjusted to 1 mL with Conjugation buffer and then combined with ANG2-P1, ANG2-P2, or ANG2-P3 peptides at molar ratios of 1:25 for ANG2-P1 and ANG2-P2, and 1:50 for ANG2-P3. The mixtures were incubated at RT for 30 min. Each mouse was injected subcutaneously with 20 μg of peptide content with an equal volume of Alum adjuvant (#77161, Thermo Scientific). The immunization schedule included a prime injection followed by three booster doses at 2, 4, and 16 weeks after the initial injection.

### Animals

All mouse experiments were conducted using C57BL/6 mice from The Jackson Laboratory, which were housed under standard conditions with continuous access to water and a maintenance diet. All mouse experiments adhered to the National Institutes of Health animal care guidelines approved by the Institutional Animal Care and Use Committees at Northwell Health and The Feinstein Institutes for Medical Research, following the NIH Guide for the Care and Use of Laboratory Animals. BMP9/10ib pups were obtained as described before^11^.

### ELISAs

The anti-BMP9 and anti-BMP10 antibody ELISAs were performed as follows: 96-well ELISA plates (Maxisorp, Nunc) were coated with 100 μL per well of recombinant BMP9 (3209-BP-010, R&D Systems) or BMP10 (2926-BP-025, R&D Systems) at 1 μg/mL in a coating buffer (15 mM K2HPO4, 25 mM KH2PO4, 0.1 M NaCl, 0.1 mM EDTA, and 7.5 mM NaN3), and incubated overnight at 4°C. The next day, the plates were rinsed three times with 0.05% Tween PBS (PBST), then blocked for 1h at RT with a 1% BSA solution in PBS. After three additional washes with PBST, serial dilutions of individual mouse serum samples were prepared, along with a reference mouse anti-BMP9 antibody (MAB3209, R&D Systems) and a reference mouse anti-BMP10 antibody (MAB2926, R&D Systems), both diluted in 1% BSA PBS. 100 μL of diluted samples were added to the wells and incubated for 2h at RT. Following three more washes, 100 μL of horseradish peroxidase (HRP)-conjugated secondary antibody (goat anti-mouse immunoglobulins, Southern Biotech; 1:500) was added and incubated for 1h at RT, followed by five washes. TMB substrate was added (#34028, Fisher Scientific), and the plate was incubated for 30 min at RT. The optical density (OD) was measured at 450 nm using a TECAN GENios Pro plate reader.

To measure anti-ANG2-P3 and anti-ANG1-P3 antibody titers in serum, 96-well plates were coated with either ANG2-P3 or ANG1-P3 peptides at 2 μg/mL. The coated plates were washed and then blocked for 1h with 5% skim milk in PBST. Serum samples were added starting at a concentration of 1:600 in 1% skim milk in 0.05% PBST, then serially diluted up to 1:307,200 and incubated for 2h at RT. After washing and adding the TMB substrate, the signal was measured. Titers were estimated by determining the IC50 value through interpolation using a sigmoidal standard curve.

### Retinal whole-mount immunofluorescence

The retinal tissues were fixed, dissected, and stained using a protocol previously described^17,18^. Briefly, freshly collected retinas were placed in a 4% paraformaldehyde solution in a 24-well plate and kept on ice for 35 min. The fixed retinas were then transferred to PBS and dissected for flat mounting. PBS was removed, and the retinas were immersed in 100% ice-cold methanol for storage or used immediately for immunofluorescence. Methanol-fixed retinas were washed in 0.3% Triton X-100 solution in PBS (PBSTX) for 15 min on a shaker at RT. The retinas were blocked in a solution containing 10% heat-inactivated goat serum, 0.2% bovine serum albumin (BSA), and PBSTX for 1h. Then, the retinas were incubated overnight on a shaker at 4°C, protected from light, with AF488-labeled isolectin GS-IB4 (#121411, ThermoFisher; 1:200) and Cy3-labeled α-smooth muscle actin (SMA; #C6198, Sigma; 1:300), diluted in the blocking solution. The following day, the retinas were washed three times for 10 min each with a washing buffer (PBSTX, 2% heat-inactivated goat serum, 0.2% BSA) and once with PBS for 5 min. Finally, the retinas were mounted on glass slides using ProLong™ Diamond Antifade Mountant (P36965, Thermo Fisher), covered with a coverslip, and sealed.

### Image acquisition and analysis

Whole-mount retinas were imaged using an EVOS M7000 fluorescent microscope. Image acquisition was performed at 2X magnification, and images were analyzed using Fiji software (ImageJ2). To identify AVMs, we used both IB4 and SMA staining. All peri-optic AVMs were quantified, regardless of their size, and the number of AVMs, as well as the total AVM surface area per retina, were recorded. Three arteries and three veins were randomly selected in each retina to measure their diameters, which were taken at 500 μm from the optic nerve center; the averages were then calculated for each retina. SMA coverage was estimated by measuring the percentage of area coverage within an 800 μm diameter circle surrounding the optic nerve. The average vessel extension was determined by measuring the length of vessels stained with IB4 from the optic nerve center to the frontal edge in each of the four petals per retina.

### Echocardiography

Mice were anesthetized with 1.8-2% isoflurane in a 1:1 mixture of oxygen and air. Hair was removed from the thoracic area, and the animals were placed on a heated platform connected to the echocardiography system (Vevo 3100, VisualSonics), which records ECG and respiratory rate. The mice’s heart rate ranged between 400 and 500 bpm. An MX-550D (40 MHz) transducer was used for image acquisition. Standard parameters were obtained using B-mode and M-mode in the “Parasternal Short Axis” (PSAX) and “Parasternal Long Axis” (PLAX) views. Multiple parameters were calculated, including cardiac wall thickness, ventricular volumes during systole and diastole, estimated left ventricle mass, ejection fraction, fractional shortening, cardiac output, and stroke volume. Pulmonary artery flow direction and velocity were assessed using Color and PW Doppler modes. Data analysis was performed using Vevo LAB 5.7.0 software.

### Biochemical serum analysis

Blood was collected into a 5 mL serum separator tube (SST, BD Vacutainer #36798). After clotting at RT for 1h, the SST tubes were centrifuged at 3500 rpm at 4°C for 15 min following the manufacturer’s instructions. The serum was then transferred to microcentrifuge tubes and centrifuged at maximum speed for 10 min at 4°C. The serum was sent to Charles River Laboratories for biochemical analysis using the Complete Clinical Chemistry panel to assess liver and kidney function.

### Statistics

All statistical analyses were performed using GraphPad Prism 10. A p-value less than 0.05 was considered statistically significant.

## RESULTS

### Immunogenicity of the ANG2-P3:CRM197 vaccine in C57BL/6 mice

Previously, our lab developed a vaccine method using minimal peptidic epitopes (down to 6 amino acids) linked to the FDA-approved carrier protein, cross-reacting material 197 (CRM 197)^16^. This approach enhances vaccine selectivity toward a specific protein or neoepitope. As potential epitopes, we first selected two conserved internal peptide sequences (ANG2-P1 and ANG2-P2) within the reported binding domain of ANG2 to Tie2^19^, which show low similarity to ANG1. Based on dose-ranging immunogenicity analyses of CRM197 vaccines and our own experience with peptide CRM197 vaccines, we administered 20 μg of vaccine (peptide content) per immunization. CRM197-coupled ANG2-P1 and ANG2-P2 failed to induce an antibody response in female C57BL/6 mice when tested 4 weeks post-prime injection (data not shown). Since CRM197 peptide vaccines are prone to respond to neoepitopes and N- or C-terminally exposed epitopes^16^, we then immunized five female C57BL/6 mice against the C-terminal last six amino acids of ANG2 (ANG2-P3 peptide, ANG2-P3:CRM197 conjugate vaccine; **Figures 1A and 1B**). ANG2-P3:CRM197 elicited a strong antibody response against ANG2-P3 (sera from vaccinated mice Vac-1 to Vac-5) when tested by ELISA at 4 weeks post-prime injection, with titers for the top three responders ranging from approximately 38,000 to 10,000 (Vac-1, Vac-2, and Vac-3), compared with controls injected with saline (Sal-1 and Sal-2) or CRM197-only (CRM-1 to CRM-5), which did not produce anti-ANG2-P3 antibodies as expected (**Figure 1C**). ANG2-P3 differs by only one amino acid from the corresponding C-terminal peptide of ANG1 (ANG1-P3; **Figure 1B**), suggesting potential cross-reactivity of ANG2-P3:CRM197 with ANG1. Indeed, we found that ANG2-P3:CRM197 elicited an immune response against ANG1-P3 peptide, but with consistently weaker titers than for ANG2-P3 (**Figures 1C, 1D**). Indeed, compared to ANG2-P3, immunogenicity toward ANG1-P3 was sixfold, twofold, and sixteenfold weaker for females Vac-1, Vac-2, and Vac-3, respectively (**Figures 1C, 1D**).

**Figure 1.**
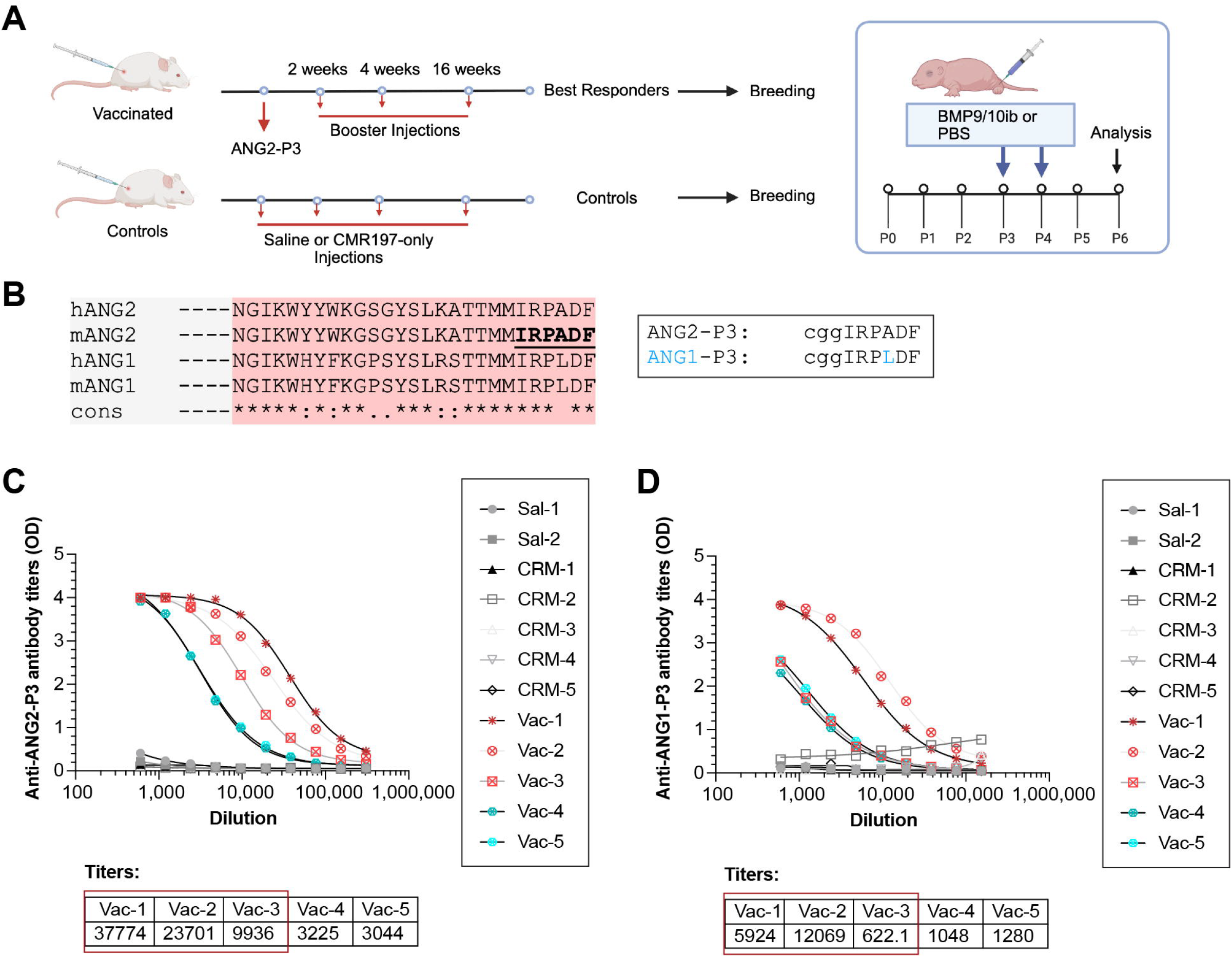
Immunogenicity of the ANG2-P3:CRM197 vaccine in C57BL/6 mice. (**A**) Schematic diagram of the vaccination schedule and generation of the BMP9/10ib model. (**B**) Protein sequence alignment of the C-terminal end of human ANG2 (hANG2), mouse ANG2 (mANG2), human ANG1 (hANG1), and mouse ANG1 (mANG1), along with the peptide sequences of ANG2-P3 and ANG1-P3. (**C and D**) Serum antibody titers against ANG2-P3 (C) and ANG1-P3 (D) in ANG2-P3:CRM197-vaccinated females (Vac-1 to Vac-5) and controls [injected with saline (Sal-1 and Sal-2) or CRM197-only (CRM-1 to CRM-5)]. The vaccinated females with the highest anti-ANG2-P3 titers were identified as “best responders” (marked with a red box). OD, optical density.

### ANG2-P3:CRM197 vaccine reduces AVM pathology in BMP9/10ib neonates

A popular model for studying HHT-type AVMs relies on postnatal development^20^ and the neutralization of BMP9 and BMP10 ligands^11,21^. Since retinal AVMs develop within the first week after birth^11^, vaccination cannot be tested in this model. Therefore, we sought to determine whether mouse neonates from vaccinated females could be exposed to anti-ANG2 antibodies and be protected against AVM development. Vaccinated females Vac-1, Vac-2, and Vac-3, along with a non-vaccinated control female, were bred with C57BL/6 males to produce litters that were injected with BMP9 and BMP10 blocking antibodies at P3 and P4, resulting in BMP9/10 immunoblocked (BMP9/10ib) pups^17^ (**Figure 1A**). ANG2-P3:CRM197 antibodies were readily detected in pups from vaccinated females, with the highest titers observed in litters from females identified as the best responders, Vac-1 and Vac-2 (**Figure 2A**). BMP9/10ib pups from vaccinated dams showed a significant decrease in retinal AVM number (**Figures 2B, 2C**) and size (**Figure 2D**). We found a negative correlation between ANG2-P3:CRM197 antibody titers and the number and size of AVMs (**Figure 2E**). Artery and vein dilation were fully normalized in BMP9/10ib retinas of pups from vaccinated dams, compared to healthy pups and BMP9/10ib pups from non-vaccinated dams (**Figs. 2F, 2G**). Furthermore, SMA coverage, which significantly increases in BMP9/10ib pups, was reduced in BMP9/10ib retinas of pups from vaccinated dams (**Figs. 2B, 2H**). However, ANG2-P3:CRM197 did not improve the retinal hyperproliferative phenotype caused by BMP9/10ib (**Fig. S1**). To ensure that the beneficial effects of the vaccination were not due to a reduction in circulating BMP9/10 antibodies, we measured anti-BMP9 and anti-BMP10 antibody levels in the sera of BMP9/10ib pups from vaccinated dams. Results showed that the sera of vaccinated pups contained the expected concentrations of anti-BMP9 and anti-BMP10 antibodies (∼60 and ∼25 μg/mL, respectively)^11^, with no significant differences between groups (**Figs. S2A, S2B**).

**Figure 2.**
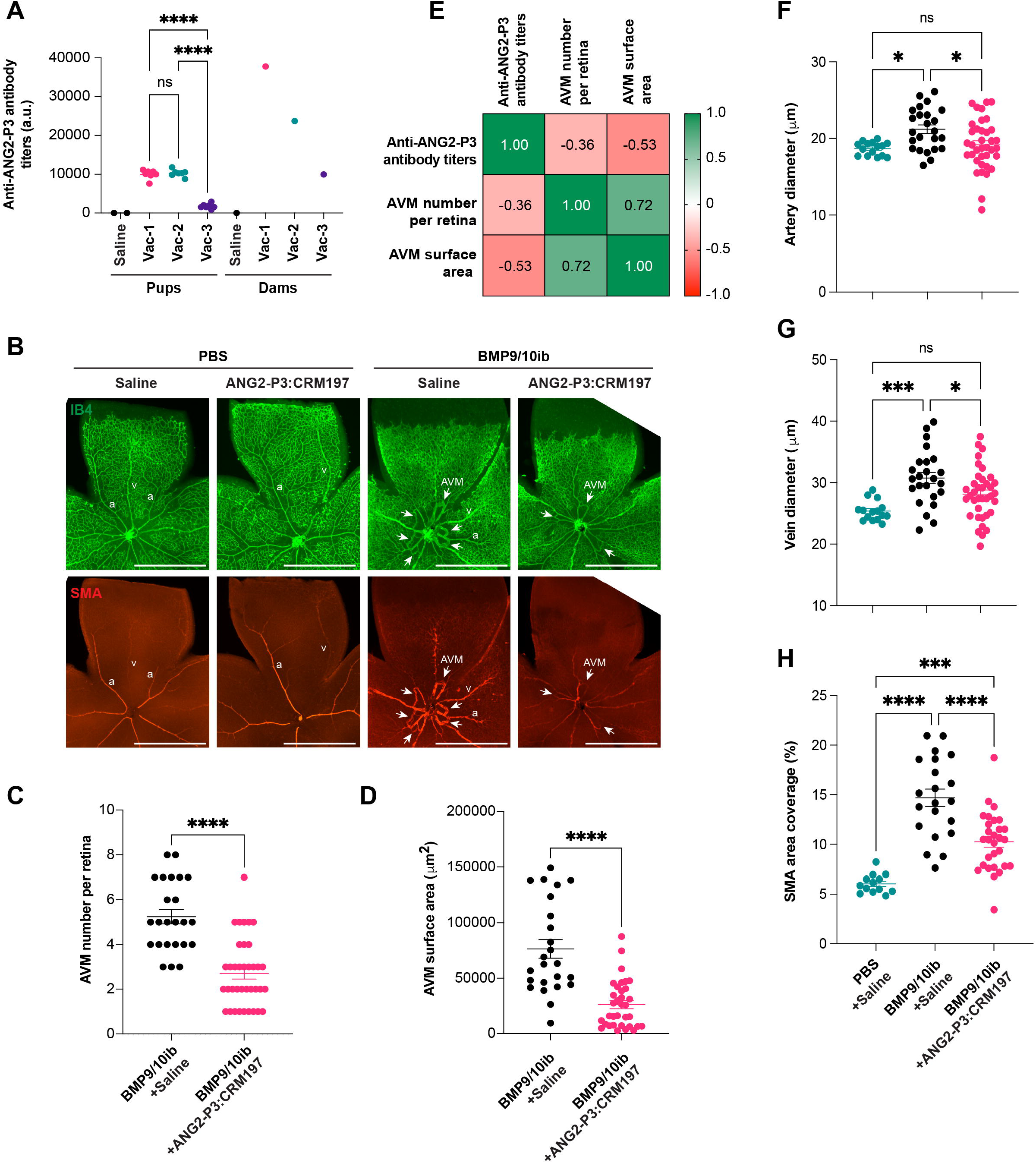
ANG2-P3:CRM197 vaccine reduces AVM pathology in BMP9/10ib neonates. (**A**) Serum antibody titers against ANG2-P3 in pups and their corresponding dams vaccinated with ANG2-P3:CRM197 (Vac-1 to Vac-3) or injected with saline (Saline). (**B**) Representative immunofluorescence staining with isolectin B4 (IB4, green) and of α-smooth muscle actin (SMA, red) in P6 retinas of pups treated with PBS or BMP9/10ib, from a dam vaccinated with ANG2-P3:CRM197 or injected with saline. a, artery; v, vein. Scale bar, 1.5 mm. (**C and D)** AVM count per retina (C) and retinal AVM surface area (D) in BMP9/10ib pups from dams vaccinated with ANG2-P3:CRM197 or injected with saline (Saline). (**E**) Spearman’s rank correlation matrix of the indicated variables. (**F-H**) retinal artery diameter (F), retinal vein diameter (G), and SMA coverage area (H) in BMP9/10ib pups from dams vaccinated with ANG2-P3:CRM197 or injected with saline (Saline). Data are shown as meanLJ±LJs.e.m.; unpaired t-test with Welch’s correction (C), Mann-Whitney test (D), and one-way ANOVA with Tukey’s multiple comparisons test (F-H). ns, not significant; ^*^P < 0.05; ^***^P ≤ 0.001; ^****^P < 0.0001.

### Mice vaccinated with ANG2-P3:CRM197 exhibit normal cardiac, liver, and kidney functions

A health check was performed on the three vaccinated females (Vac-1, Vac-2, and Vac-3) as well as on three age-matched non-vaccinated females who received saline injections. An echocardiogram was performed to assess heart and lung health, and a complete blood count was conducted for liver and kidney function testing, one year after the final booster, and after all offspring had been collected. Ultrasound imaging of the heart, using B- and M-modes, was analyzed for multiple parameters (**Figure S3**). Pulmonary acceleration time (PAT) and the ratio of PAT to pulmonary ejection time (PAT/PET) assessed via B-mode serve as indicators of pulmonary arterial hypertension and right ventricular health. No significant differences were observed between vaccinated and non-vaccinated groups. The assessment also included the left ventricle area, a crucial component of the circulation responsible for pumping oxygenated blood to nearly all organs. Results showed no significant difference in the left ventricle area between the two groups. We used M-mode to measure several parameters on the parasternal short-axis (PSAX) and long-axis (PLAX) views, including heart rate, which showed no significant difference between groups. Cardiac output, estimating the total blood volume the heart pumps per minute, and ejection fraction, representing the percentage of blood ejected per heartbeat, were also measured. Fractions of shortening, indicating the left ventricle’s contractile effectiveness, and left ventricular mass were examined, too; none showed significant differences.

To assess the impact on liver and kidney functions, we analyzed metabolites and enzymes such as aspartate aminotransferase (AST), alanine aminotransferase (ALT), alkaline phosphatase (ALP), gamma-glutamyl transferase (GGT), creatine kinase (CK), creatinine, total bilirubin, urea nitrogen, glucose, total cholesterol, triglycerides, total protein, albumin, globulin, albumin/globulin ratio, calcium, phosphorus, sodium, potassium, and chloride. No significant differences were found in any of these parameters between vaccinated and non-vaccinated mice (**Figure S4**).

## DISCUSSION

ANG2-neutralizing antibodies have demonstrated promising results by targeting ANG2 elevation and decreasing vascular pathology in different HHT and AVM models^12,13,22^. Therapeutic monoclonal antibodies have become increasingly important due to their targeted biological effects and ability to treat a variety of conditions, including autoimmune diseases and cancer^23^. However, monoclonal antibody therapies have some drawbacks, including high costs, frequent dosing, and the risk of developing antibody resistance^24,25^. Several peptide-based synthetic vaccines (or peptide vaccines) are now FDA-approved, especially for viral diseases and cancer^26^. Vaccinal approaches activate both humoral and cellular immune responses, providing a broader and more durable effect than passive immunization; this long-lasting benefit may be an essential alternative for addressing persistent ANG2 elevations, as vascular complications pose a lifelong challenge for HHT patients.

Previously, our lab developed a selective peptide vaccine that uses minimal peptidic epitopes linked to CRM197. This method enabled the creation of a vaccine that targets specifically the N-terminally truncated pyroglutamate-3 pathological form of the amyloid-β protein, a key component of amyloid plaques in the brains of Alzheimer’s disease patients^16^. Here, we describe the development and initial characterization of a CRM197-conjugated ANG2 peptide vaccine. We report the identification of the ANG2-P3:CRM197 vaccine and demonstrate that it significantly reduces AVM number and size, as well as normalizes arterial and venous diameters in the retina of the BMP9/10ib model of HHT. Vaccinated mice exhibit normal cardiac, liver, and kidney functions, indicating that the ANG2-P3:CRM197 vaccine does not cause toxicity under the tested conditions one year post-immunization. Further research will be necessary to assess the safety of ANG2 neutralization with ANG2-P3:CRM197 in more challenging scenarios, such as those involving angiogenic responses during inflammation or physical trauma.

The role of ANG2 in HHT pathogenesis remains unclear. ANG2 is an endothelial tip cell marker that regulates angiogenesis and belongs to the angiopoietin growth factor family^27,28^. ANG1, another member of the angiopoietin growth factor family, is a potent agonist of Tie2, a tyrosine kinase receptor promoting EC stabilization and quiescence. ANG2 can act as either an agonist or antagonist of Tie2, depending on the physiological context, and is often associated with EC destabilization and vascular activation^10^. Recently, the Meadows lab, in collaboration with our team, provided evidence that the increase of ANG2 triggered by ALK1 LoF interferes with ANG1/Tie2 signaling, suggesting that ANG2, in this context, functions as an antagonist of Tie2. In the postnatal retinal model of angiogenesis, we demonstrate that the ANG2-P3:CRM197 vaccine reduces the number and size of AVMs and normalizes vessel dilation; however, it does not prevent the hypervascularization observed in this model. Hypervascularization in the retina of HHT models mainly occurs at the front of the developing vascular bed, where tip cells are found in higher numbers compared to normal retinas^29^. Future work should explore why our vaccine did not improve the hypervascularization phenotype in this model, despite ANG2’s key role in angiogenesis. It is possible that other tip cell-secreted regulators, such as apelin and Esm1, which are also elevated in HHT models^13^, contribute to the increased vascularization in HHT pup retinas and may also need to be targeted for neutralization to effectively interfere with this process.

In summary, we suggest that ANG2 neutralization and reduction of vascular pathology severity in HHT mice can be achieved with a safe ANG2 vaccine. This method offers opportunities for sustained, long-lasting anti-AVM protection in HHT, further supporting the notion that targeting ANG2 is a promising strategy for treating HHT.

## ACKNOWLEDGEMENTS

We thank Dr. Stryder M. Meadows (Tulane University) for providing insightful comments on this manuscript. This work was supported by National Institutes of Health grants R01HL139778, R01HL150040, and R01HL163196. Schematics were created with Biorender.

## AUTHORSHIP CONTRIBUTIONS

Conceptualization: PM, SR, SQ

Methodology and investigation: SQ, HZ, ZW, CI, SR

Writing: PM, SQ

