## Supplementary Figures S1-S4 for "An angiopoietin-2 vaccine improves arteriovenous malformation pathology in hereditary hemorrhagic telangiectasia mice"

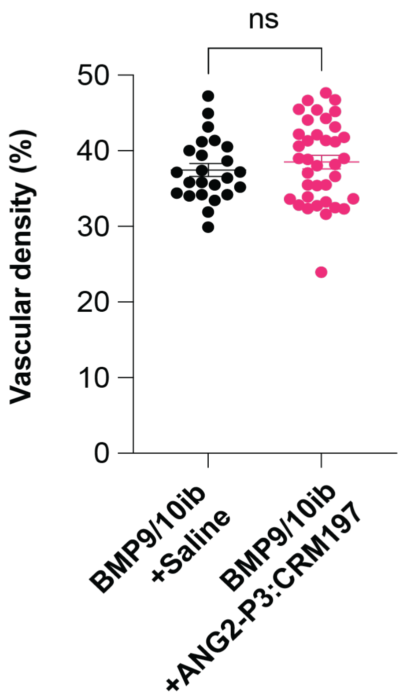


**Figure S1: Effect of the ANG2-P3:CRM197 vaccine on retinal hypervascularization in BMP9/10ib neonates.** Average vascular density measurement in the plexus between arteries and veins in BMP9/10ib pups from dams vaccinated with ANG2-P3:CRM197 or injected with saline (Saline). Data are shown as mean ± s.e.m.; unpaired t-test with Welch's correction. ns, not significant.

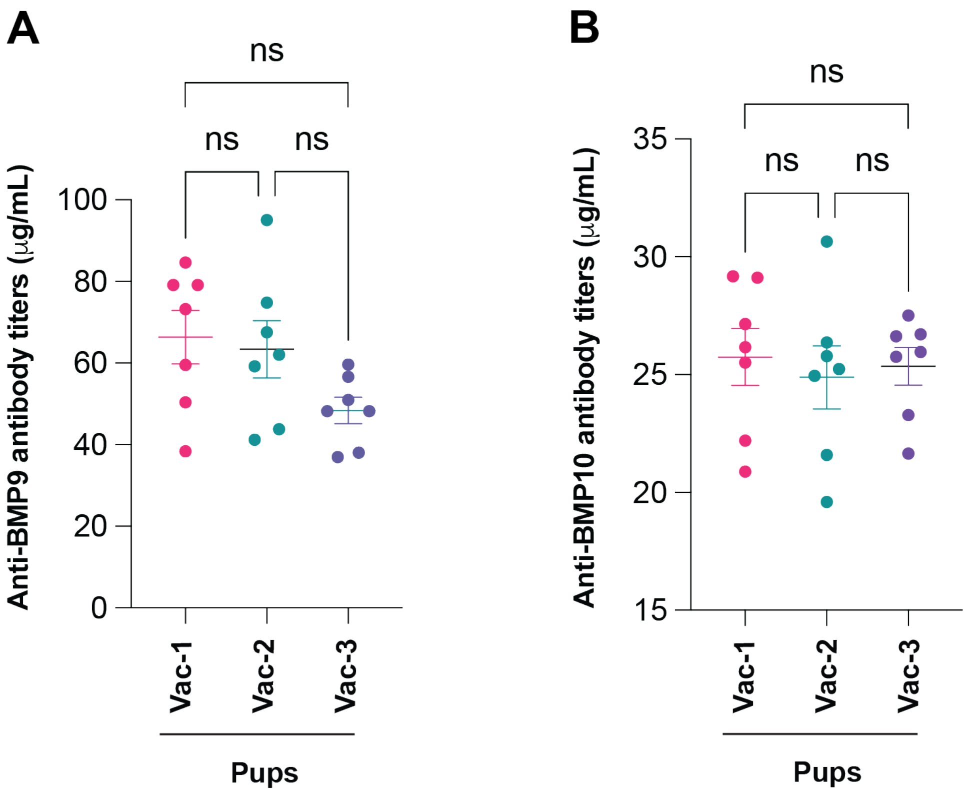


**Figure S2: Circulating anti-BMP9 and anti-BMP10 antibody concentrations in BMP9/10ib neonates from ANG2-P3:CRM197-vaccinated dams. (A and B)** Serum anti-BMP9 (A) and anti-BMP10 (B) antibody concentrations in BMP9/10ib neonates from the three best responders of the ANG2-P3:CRM197-vaccinated dams (Vac-1 to Vac-3). Data are shown as mean ± s.e.m. (n=7); one-way ANOVA with Tukey's multiple comparisons test. ns, not significant.

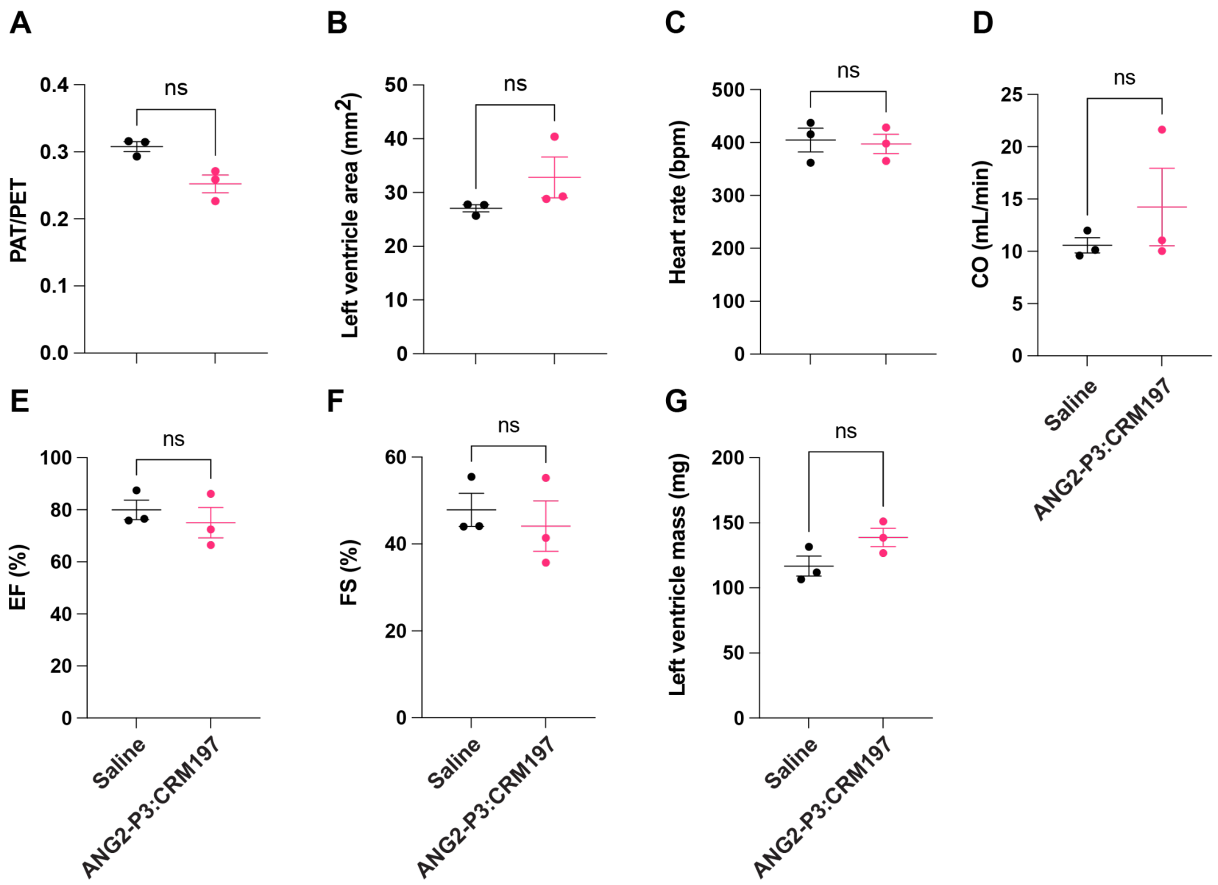


**Figure S3: Echocardiography measurements in ANG2-P3:CRM197-vaccinated females.** (**A-G**) Echocardiographic data in ANG2-P3:CRM197-vaccinated and control (Saline) females 12 months post vaccination: (A) Pulmonary acceleration time (PAT)/pulmonary ejection time (PET) ratio and (B) left ventricle area measured in PSLAX B-mode (C) heart rate, (D) cardiac output (CO), (E) ejection fraction (EF), (F) fractional shortening (FS), and (G) left ventricle mass measured in PSLAX M-mode. Data represent individual mice and mean ± s.e.m. (n=3); Mann-Whitney test. ns, not significant.

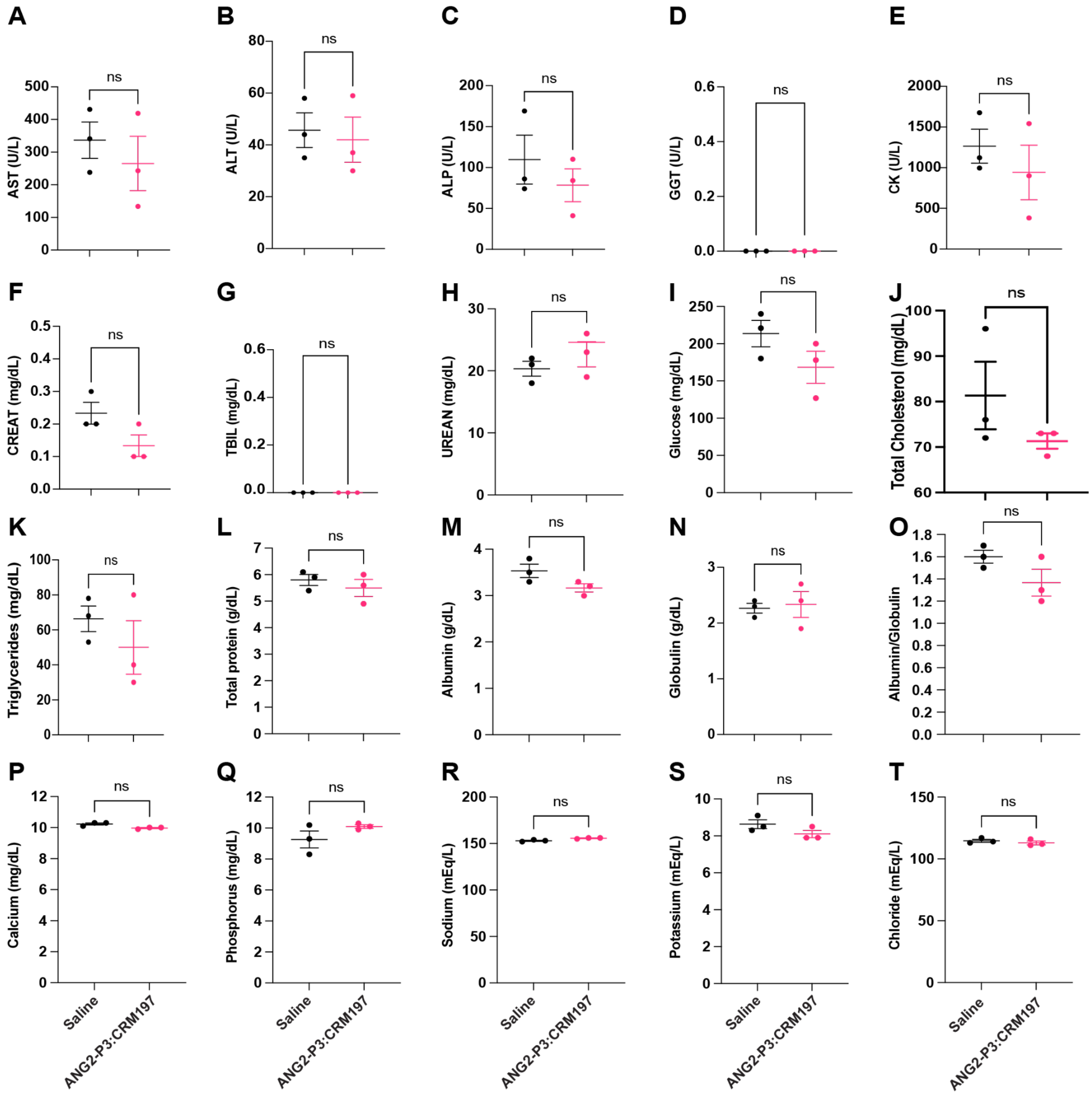


**Figure S4: General liver and kidney function assessment in ANG2-P3:CRM197-vaccinated females. (A-T)** The following serum measurements were taken in ANG2-P3:CRM197-vaccinated and control (Saline) females 12 months post vaccination were made in the serum: Aspartate aminotransferase (AST, A), alanine aminotransferase (ALT, B), alkaline phosphatase (ALP, C), gamma-glutamyl transferase (GGT, D), creatine kinase (CK, E), creatine (CREAT, F), total bilirubin (TBIL, G), urea nitrogen (UREAN, H), glucose (I), total cholesterol (J), triglycerides (K), total protein (L), albumin (M), globulin (N), albumin/globulin ratio (O), total calcium (P), phosphorus (Q), sodium (R), potassium (S), and chloride (T). Data represent individual mice and mean ± SEM; (n=3) for each group; Mann-Whitney test; ns, not significant.
